# Improved protein binder design using beta-pairing targeted RFdiffusion

**DOI:** 10.1101/2024.10.11.617496

**Authors:** Isaac Sappington, Martin Toul, David S. Lee, Stephanie A. Robinson, Inna Goreshnik, Clara McCurdy, Tung Ching Chan, Nic Buchholz, Buwei Huang, Dionne Vafeados, Mariana Garcia-Sanchez, Nicole Roullier, Matthias Glögl, Christopher J. Kim, Joseph L. Watson, Susana Vázquez Torres, Koen H. G. Verschueren, Kenneth Verstraete, Cynthia S. Hinck, Melisa Benard-Valle, Brian Coventry, Jeremiah Nelson Sims, Green Ahn, Xinru Wang, Andrew P. Hinck, Timothy P. Jenkins, Hannele Ruohola-Baker, Steven M. Banik, Savvas N. Savvides, David Baker

## Abstract

Despite recent advances in the computational design of protein binders, designing proteins that bind with high affinity to polar protein targets remains an outstanding problem. Here we show that RFdiffusion can be conditioned to efficiently generate protein scaffolds that form geometrically matched extended beta-sheets with target protein edge beta-strands in which polar groups on the target are nearly perfectly complemented with hydrogen bonding groups on the design. We use this approach to design binders against a set of therapeutically relevant polar targets (KIT, PDGFRɑ, ALK-2, ALK-3, FCRL5, and NRP1) and find that beta-strand-targeted design yields higher affinities and success rates than unconditioned RFdiffusion. All by all binding experiments show that the designs have affinities ranging from 137 pM to mid nM for their targets and essentially no off target binding despite the sharing of beta-strand interactions, likely reflecting the precise customization of interacting beta-strand geometry and additional designed binder-target interactions. A co-crystal structure of one such design in complex with the KIT receptor is nearly identical to the computational design model confirming the accuracy of the design approach. The ability to robustly generate binders displaying high affinity and specificity to polar interaction surfaces with exposed beta-strands considerably increases the range and capabilities of computational binder design.

## Introduction

There has been considerable recent progress in de novo protein binder design^1–6^. Both energy based^1^ and deep learning methods, such as RFdiffusion^2^, now enable design of protein binders given only the structure of the target of interest and (optionally) the specification of region of the target surface to bind^3^. Despite this progress, design of high affinity binders to polar regions of a target surface remains challenging since the exposed hydrogen bond donors and acceptors must be complemented with precisely positioned acceptors and donors on the designed binder to compensate for the loss of interactions with water. While RFdiffusion excels at generating backbones that are shape complementary to the targeted region of a protein surface, these solutions do not always provide detailed complementarity of hydrogen bonding donors and acceptors^7^. Many therapeutically relevant target proteins have beta-sheets with unpaired and exposed beta-strands; these often have non-canonical structures to reduce the tendency for aggregation^8,9^. Beta-strand targeted binders have been generated using pre-deep learning Rosetta methodology^10^, but this approach has limited ability to match the diversity of edge beta-strand geometries. A generalizable deep learning method for designing beta-strand pairing based binders to complement arbitrary target beta-strand twists^11^, bends^12^, bulges^13^, and other irregular features^14,15^ could improve the design of binders to polar target surfaces.

We reasoned that RFdiffusion generation of binder backbone mediated hydrogen bonding interfaces with the target could yield designs with polar groups nearly perfectly complementing those on the target. We set out to develop a general approach for guiding RFdiffusion denoising-diffusion trajectories towards such beta-strand centric interfaces, and we hoped to experimentally validate this new binder design approach.

### beta-strand pairing conditioning improves protein interface design quality

During training of RFdiffusion, a subset of training examples were provided with secondary structure and secondary structure block-adjacency (SS/ADJ) information; this was found to enable the conditioning of the model towards user defined protein monomer folds^2^. We explored whether providing interface conditioning information indicating a binder beta-strand pairing with a target edge beta-strand could yield designable strand pairing complexes.

RFdiffusion takes as input desired residue secondary structures as well as a N by N block-adjacency matrix (N is the length of the designed protein) which specifies desired adjacencies between secondary structure blocks. We developed a simple algorithm for mapping user specified parameters – the length of the designed interacting beta-strand (L) and the identity of the target beta-strand to be bound (T) – onto tensors for conditioning the binder-target interface. When provided this conditioning information, RFdiffusion generates binders assigning a random set of L consecutive binder residues to form a beta-strand pair with the target strand T, while the remainder of the binder output residues are not explicitly assigned a secondary structure or target interaction. We explored using these to condition RFdiffusion and found that outputs from conditioned beta-strand pairing runs indeed contained the user specified beta-strand interfaces (Figure 1a-b, Extended data figure 1).

**Figure 1.**
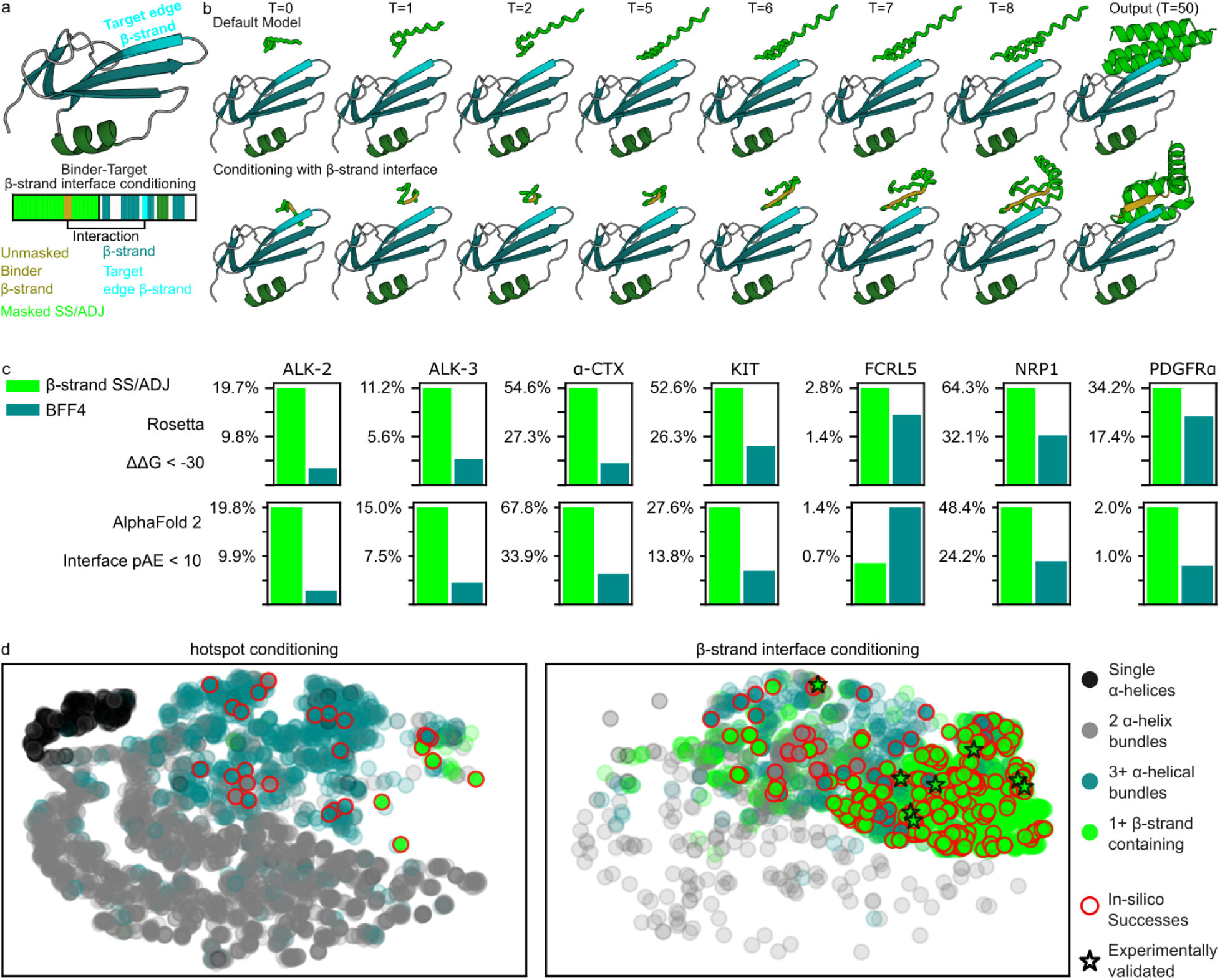
Design of beta-strand pairing binders. **(a)** Representation of beta-strand interface conditioning information provided as a tensor to RFdiffusion to generate beta-strand pairing binders. **(b)** Example RFdiffusion denoising trajectories without (top) and with (bottom) beta-strand interface conditioning. Conditioning indicates that part of the binder scaffold should be a beta-strand (gold) that contacts the indicated target edge strand (cyan). This information influences the denoising in very early trajectory timesteps (t), with the tertiary fold determined within 5 timesteps and the final output at t=50. **(c)** Binder design success rates. **(d)** Structural clustering of RFdiffusion output binder scaffolds using hotspot conditioning (left) or beta-strand interface conditioning (right) t-SNE transforms of all-by-all pairwise RMSD data among binder scaffolds across all targets are plotted, with close proximity of points representing structural similarity. Output fold secondary structures are classified by color as indicated in the legend. Bold bordered data points indicate In silico successes (red circles) and experimentally validated binders (black stars).

We supplied RFdiffusion with beta-strand interface conditioning tensors for larger scale binder design campaigns against protein targets containing edge beta-strands sufficiently exposed for binder access. To evaluate the generality of the approach, we selected targets that span a range of edge beta-strand geometries. The seven selected targets also have considerable therapeutic relevance. Activin receptor-like kinases 2 and 3 (ALK-2 and ALK-3) are both Type I bone morphogenetic protein receptors that regulate growth of bone, vasculature, hair follicles, enamel as well as wound healing and tumor suppression in various soft tissues^16–20^. Platelet-derived growth factor receptor ɑ (PDGFRɑ) and Mast/stem-cell factor receptor (also known as KIT, SCFR, CD117) are both type III receptor tyrosine kinases that play roles in cardiomyocyte proliferation and heart tissue regeneration after myocardial infarction^21–29^.

FCRL5 functions in critical signaling pathways for B-cell activation^30^. NRP1 is a coreceptor for various growth factor signaling pathways (TGF-1, EGF, VEGF, PI3K, HGF, and PDGF)^31^, a viral entry factor^32^, and plays a role in the of RAS/MAPK signaling in various cancers^33,34^. ɑ-Cobratoxin (ɑ-CTX), the lone non-receptor target, is a major paralytic component of cobra venom^35^ that acts by blocking muscle and nerve acetylcholine signaling. All except ɑ-CTX are therapeutic targets for different cancers^36–48^. Designed protein binders against these targets could be useful for antagonizing native signaling, targeting drug conjugates and other therapeutics to tumors, designing novel agonists^49–51^ and targeted receptor degradation^52^, inhibitors^1^, triggering cargo endocytosis^52^, or target therapeutics towards particular cell types^53^.

The set includes single beta-sheet targets with highly exposed edge beta-strands (ɑ-CTX, FCRL5, NRP1) and somewhat occluded edge beta-strands (ALK-2 and ALK-3), as well as immunoglobulin (Ig) fold beta-sandwich domains (FCRL5, KIT, and PDGFRɑ). All had been found to be challenging for binder design using standard hotspot-only conditioned RFdiffusion–either in silico design calculations had failed to generate designs predicted with high confidence to bind to the target (PDGFRɑ, FCRL5, ɑ-CTX, NRP1) or experimental testing had failed to yield binders (ALK-2, ALK-3, KIT). Because no experimentally determined structure exists for FCRL5, we designed binders against the AlphaFold 2 model^54,55^. To provide a stringent test of design specificity, the set includes some structurally related targets: ALK-2 and ALK-3 are members of the TGF-beta superfamily^56^, and KIT and PDGFRɑ are type-III receptor-tyrosine kinases^57^ (Extended data figure 2).

**Figure 2.**
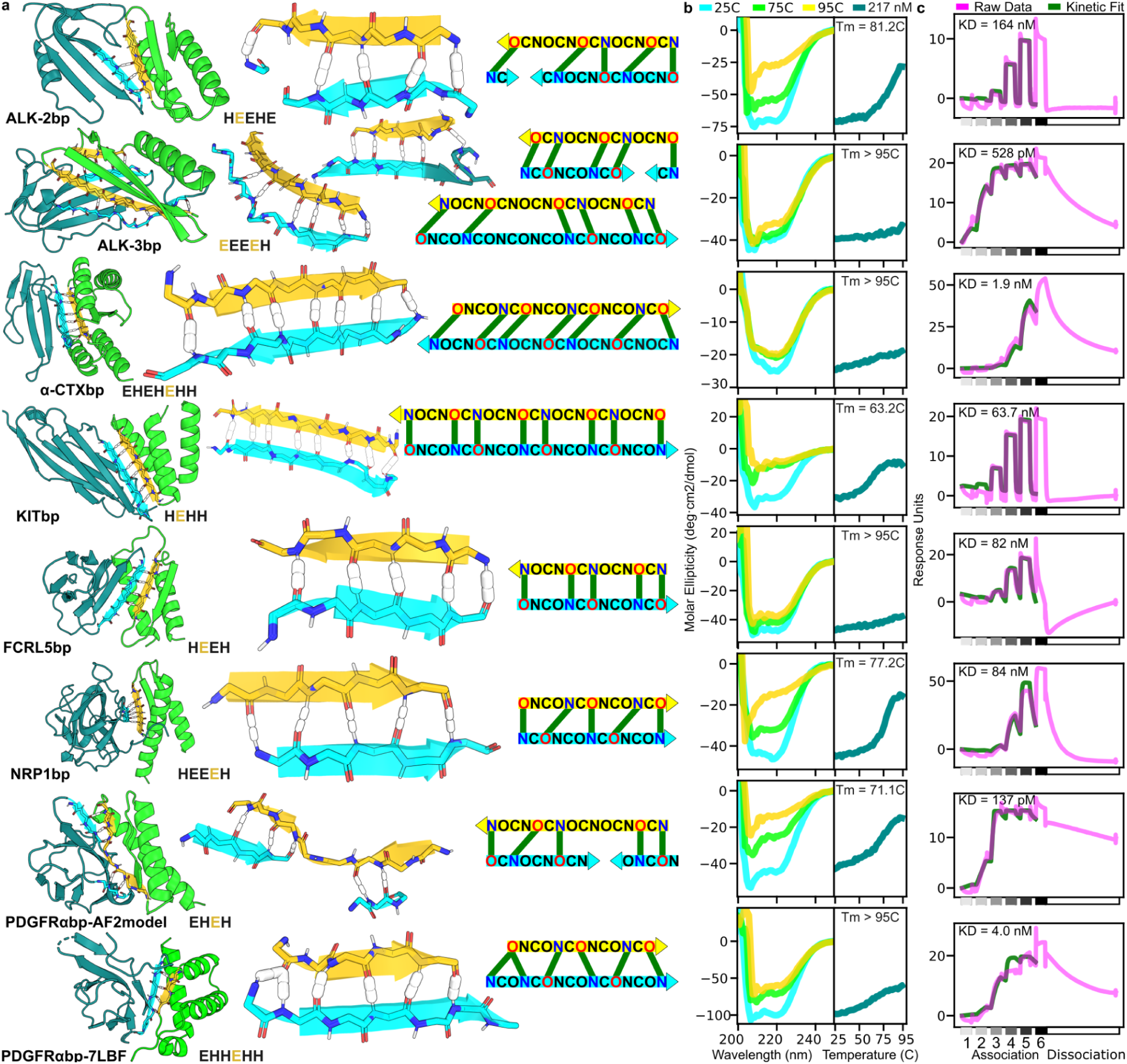
Binder design models and biophysical characterization. **(a)** Design models of binding complexes for each target. Left: Entire complex design models with binders indicated in green/gold and targets in teal/cyan. Beta-strand interface conditioned scaffold design yielded binders with interfacial beta-strands (gold) forming strand-pairing hydrogen bonding interactions with target edge beta-strands (cyan). Middle: Close up view of design model backbone hydrogen-bond interactions with putative hydrogen bonds shown in white. Right: schematic representation of strand pairing interactions showcase the diversity of sequence-independent beta-strand pairing interactions. **(b)** Circular dichroism thermal melts. Full spectrum analyses (left) performed at 25C (cyan), 75C (green), and 95C (gold) assess the overall binder fold at these three temperatures, while single wavelength thermal melts (right) were measured at 217 nM to calculate binder Tm values. **(c)** SPR measurement (pink) of binding kinetics at 600pM, 4nM, 30nM, 200nM, 1.5μM, and 10μM (association phases 1-6 on the X-axis). Fits for *K*_d_ determination (green) excluded the 10μM data excluded due to signal aberrations at this high binder concentration.

We compared the diversity and quality of binders designed using our interface conditioning method to designs generated using the standard RFdiffusion target site “hotspot” directed approach. Following the generation of 500 scaffolds using each diffusion approach, ten candidate binder sequences were generated for each scaffold by ProteinMPNN^58^ and each binder candidate complex was predicted with AlphaFold 2^59^. Across all seven targets, beta-strand interface conditioning yielded design models with improved in silico binding metrics^6^ (AlphaFold2 interface predicted aligned error (pAE) and Rosetta ΔΔG, Figure 1b), with over 9.2% of beta-strand interface conditioned designs meeting reasonable quality metrics, in contrast to 0.98% success rates by RFdiffusion conditioned on target hotspots alone.

beta-strand interface conditioning yielded in silico design successes for all targets, whereas hotspot conditioning alone did not yield any in silico successes for two targets (ALK-3 and FCRL5). beta-strand interface conditioning also yielded more globular binder designs with 88.7% of output scaffolds meeting radius of gyration criteria and only 25.5% of designs from the other methods meeting the same criteria (Extended data figure 3). beta-strand interface conditioning yielded outputs that covered a distinct protein structure space compared to other methods, as quantified by all-by-all pairwise RMSD between aligned binder scaffolds (Figure 1c), with as expected a higher fraction of beta-sheet containing binders and a decrease in ɑ-helical bundle outputs (Figure 1d).

**Figure 3.**
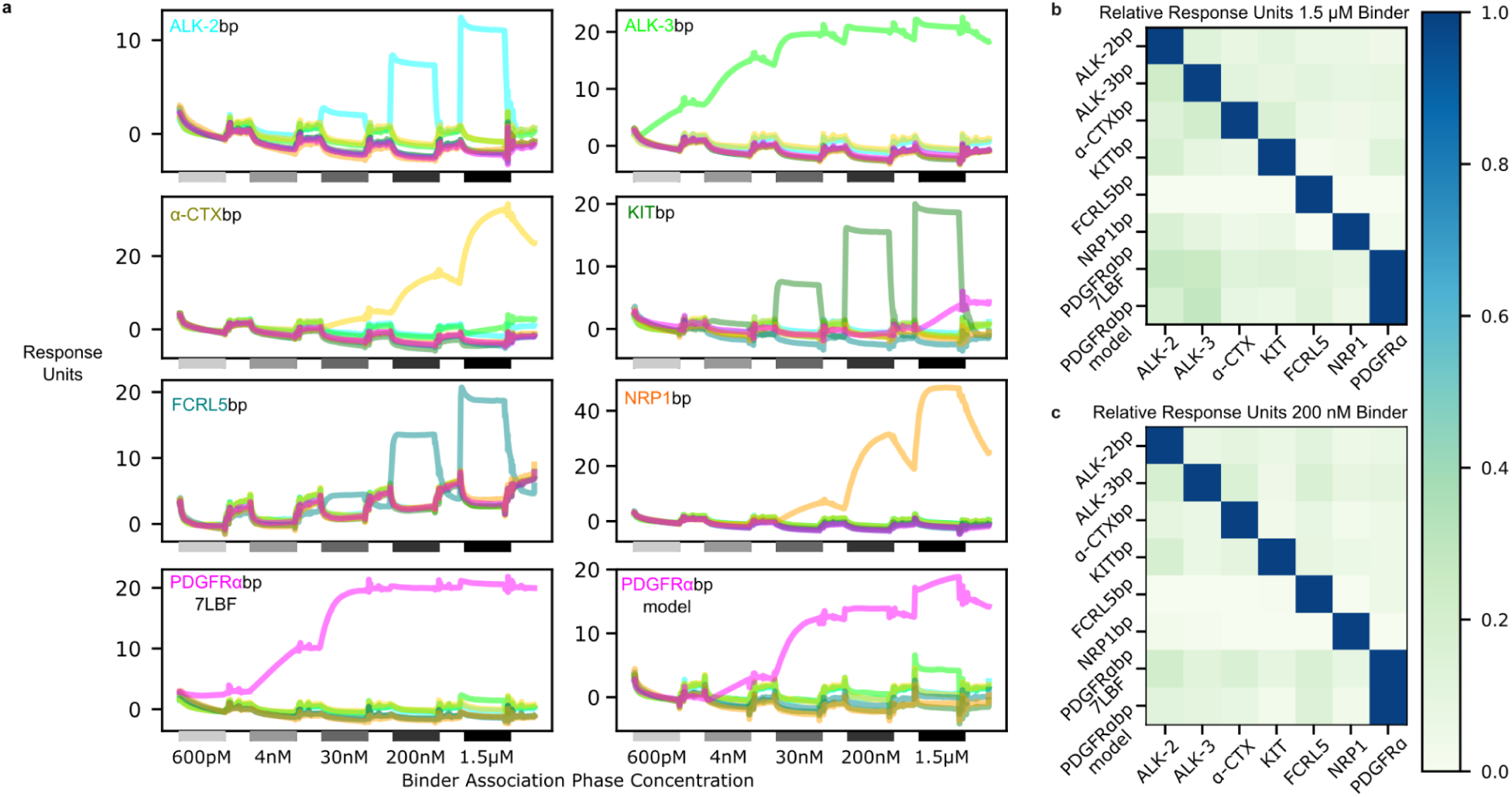
Designed binders are highly specific for their targets. **(a)** Designed binder SPR response traces against all targets. For each binding protein, immobilized cognate target protein yielded the strongest SPR binding response at various binder concentrations (7-fold increasing binder concentrations ranging from 600pM to 1.5μM, from left to right in each trace). **(b,c)** Average response units for the **(b)** 1.5μM and **(c)** 200nM binder concentrations. Grid colors indicate response units relative to the maximum response for each binder.

For each target, sets of 100-10^5^ designs were evaluated experimentally. The sets were generated using beta-strand targeted RFdiffusion and with standard hotspot conditioned RFdiffusion, followed by ProteinMPNN, and selection based on AlphaFold 2 and Rosetta metrics; partial diffusion scaffold optimization^4^ and additional sequence sampling was carried out to generate designs with improved metrics in several cases. Design sets were first screened using yeast surface display. Significantly more binders were obtained with the strand directed approach than with standard RFdiffusion at this selection stage (Extended data figure 4). The tightest binders were expressed in *E. coli* and purified (Extended data figure 5). SPR revealed mid-to sub-nanomolar affinity binders for our targets (Figure 2a-c). In the following sections we describe the active designs for each target in turn.

**Figure 4.**
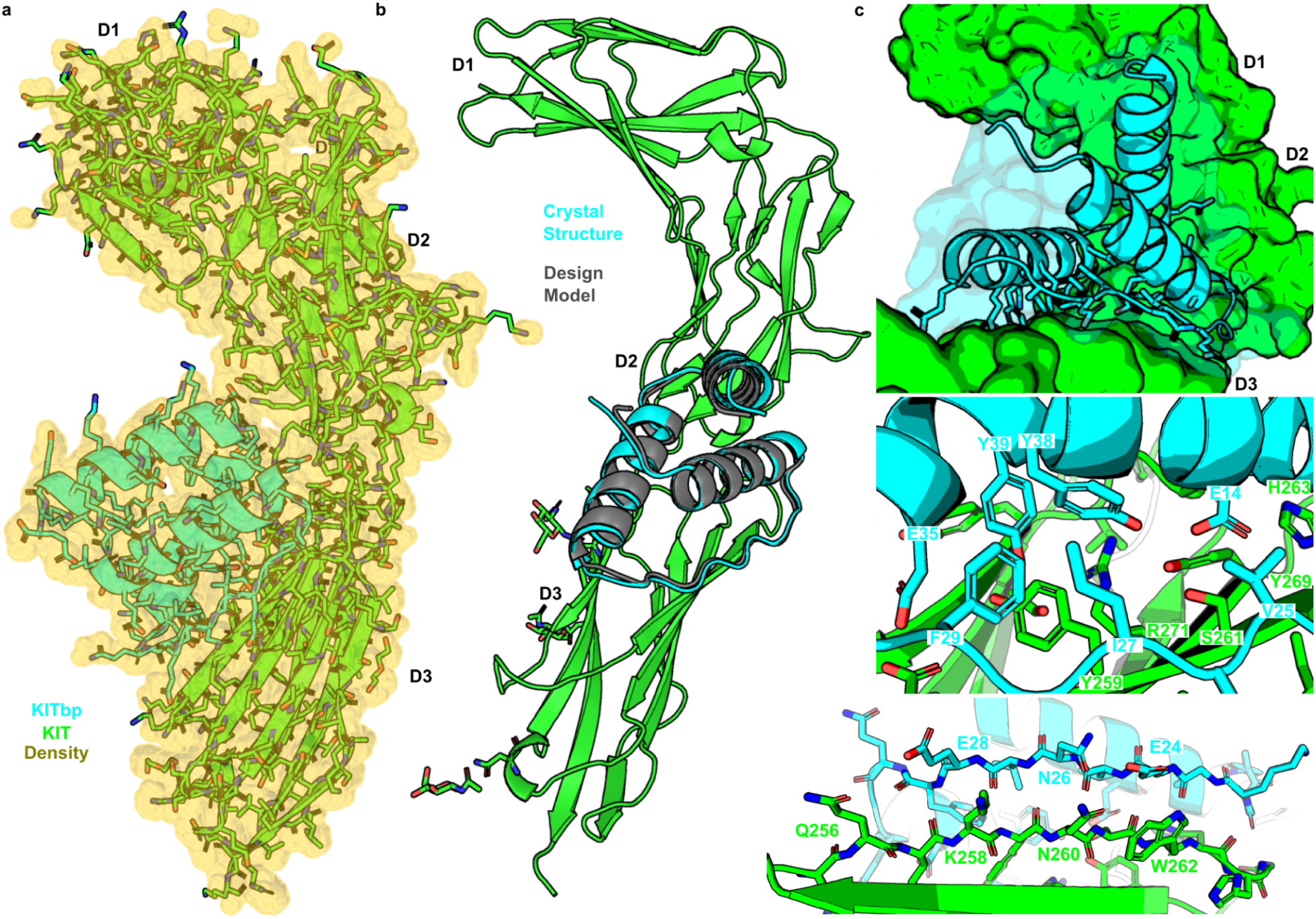
Structural analysis of the KITbp-KIT complex. **(a)** Crystal structure (PDB ID: 9H71) of KITbp (cyan) in complex with KIT (green) and 2Fo-Fc electron density contoured at +1.0 RMSD (gold). **(b)** The KITbp crystal structure superimposes on the binder design model (grey) with subangstrom backbone atom RMSD, and 1.8 Å all-atom RMSD. **(c)** Close-up views of the binder interface reveal high shape complementarity of the binder-target complex (top). The binder interface consists of hydrophobic and polar interactions between binder core-boundary residues and KIT receptor domain 2 core-boundary residues (middle). The binder forms an extensive hydrogen-bond beta-pairing interaction with the targeted domain 2 edge strand (bottom). There are multiple side-chain interactions between binder and KIT tyrosine residues (Y38 and Y39 in the binder; Y259 and Y269 in KIT) in the core-boundary interaction, and between binder glutamate residues (E14, E28, E24, E35) and KIT hydrogen bond donors (Q256, K258, N260, W262, H263, R271). The binder also makes electrostatic interactions with both the opposite domain 2 edge strand and KIT domain 3.

**Figure 5.**
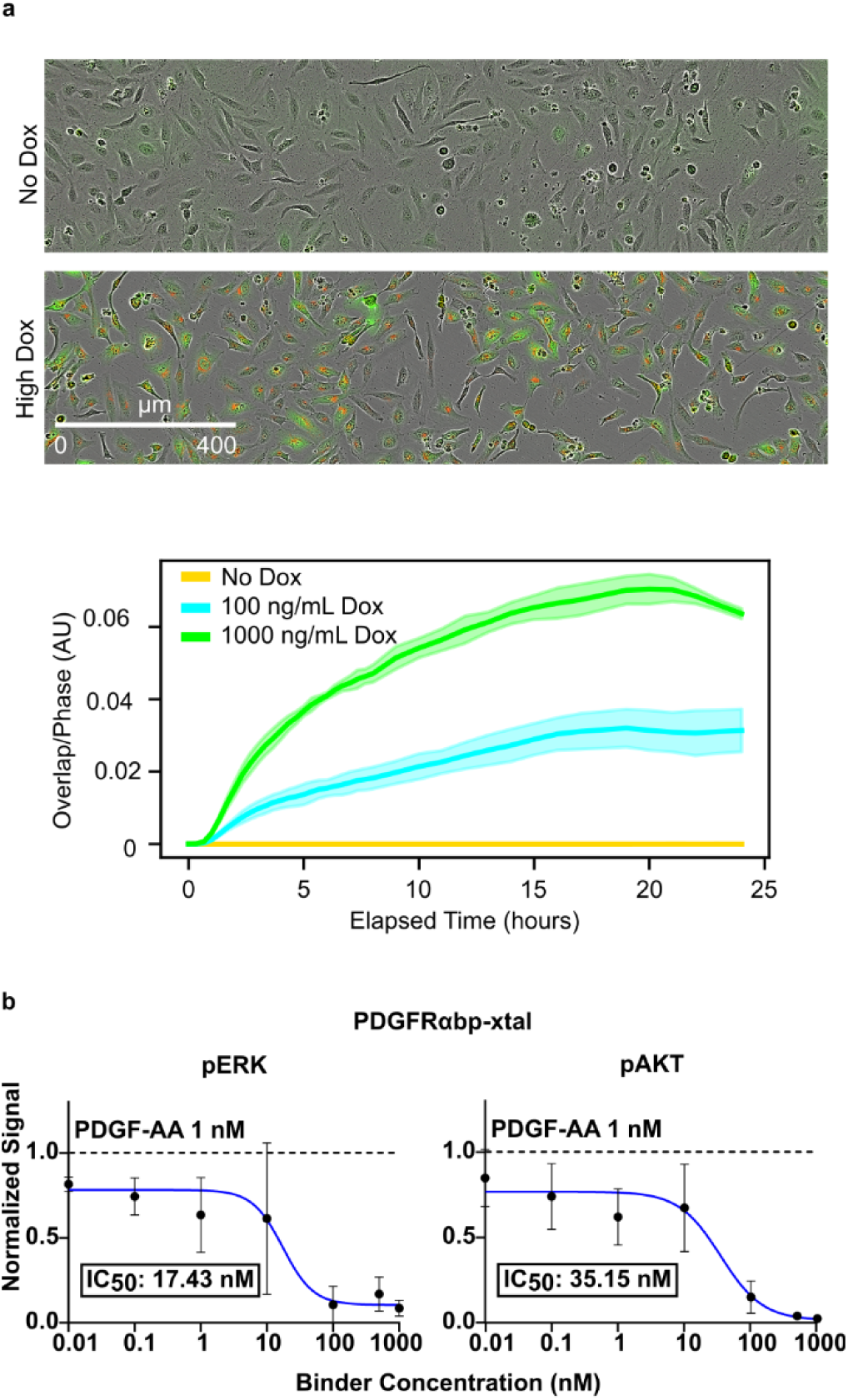
Functional activity of FCRL5 and PDGFRα binding proteins. **(a)** HeLa cells engineered to express FCRL5 receptor in a Dox-inducible manner were treated with 50nM neutravidin-labeled pHrodo DeepRed complexed FCRL5bp, and live cell imaging (top panels) was used to measure overlap of GFP (FCRL5-expressing cells) and pHrodo Red fluorescence (internalized FCRL5bp). 1000 ng/mL doxycycline results in strong GFP expression and intracellular pH activated pHrodo Red fluorescence. Internalization (measured by overlap of red and green fluorescence) reached steady state in 18 hours (bottom panel). **(b)** Western blot analysis of PDGFRα inhibition in Chinese Hamster Ovary cells engineered to overexpress PDGFRα. Levels of phosphorylated PDGFRα, Erk, and Akt were measured by immunoblots with fluorescent antibodies. Signals were normalized by the fluorescent signal of an antibody against the constitutively expressed housekeeping proteins S6 or actin.

### TGF-beta superfamily Targets: ALK-2 and ALK-3

ALK-2 and ALK-3 are two of four known Type I Bone Morphogenetic Protein Receptors (BMPRI). Targeting BMPR therapeutics towards specific tissues and pathologies could have considerable utility, but achieving binding specificity for ALK-2 versus ALK-3 presents a difficult challenge due to their structural homology. The ALK-2 and ALK-3 ectodomains have 30% sequence identity and high structural similarity (Extended data figure 2). On both targets, there is an edge beta-strand with five consecutive residues with backbone atoms in very similar atomic positioning (RMSD 0.07), and each strand is partially occluded by an N-terminal coil. Prior to this study, multiple design campaigns – including recent campaigns utilizing RFdiffusion, ProteinMPNN, and AlphaFold 2 – had failed to yield binders.

We conditioned RFdiffusion on generating beta-strand pairings to these five edge beta-strand residues that were not occluded by the N-terminal coil. We identified binding proteins, ALK-2bp and ALK-3bp, with *K*_d_ values of 164 nM and 528 pM, respectively (Figure 2c) by surface plasmon resonance (SPR). While both binders were designed to strand pair with the homologous target edge beta-strand, the actual binder interaction footprints were quite distinct, with ALK-2bp forming a parallel beta-strand interaction with the target beta-strand and ALK-3bp pairing to the same strand in an antiparallel manner. ALK-3bp makes additional strand-pairing like contacts with the C-terminal coil of ALK-3 in a non-canonical hydrogen bonding pattern, with the binder beta-strand curvature nearly perfectly complementing a bulge in the coil motif; these additional interactions may account for the four orders of magnitude greater binding affinity. This additional hydrogen bonding network with the C-terminal coil was not prespecified; RFdiffusion simply generated the secondary binder beta-strand given the context of the initial beta-strand motif specified by interface conditioning.

### Single Exposed Edge beta-strand: ɑ-CTX, FCRL5, and NRP1

Three of the targets have more exposed edge beta-strands. The acetylcholine receptor antagonist ɑ-CTX is a three-finger toxin that consists of a single beta-sheet with extended loops that bind a hydrophobic receptor pocket. We aimed to design a beta-strand pairing binder that would sterically interfere with this interaction. FCRL5 weakly binds IgG to modulate B-cell activation. While the exact IgG binding site is unknown^60^, the receptor consists of several Ig domains. In the AlphaFold2 model, the N-terminal Ig domain contains one three stranded sheet and one five stranded sheet, leaving an exposed beta-strand available to target during design. Finally, we targeted the discoidin domain of NRP1, which contains a highly twisted beta-sheet with a potential beta-strand binding site where the edge beta-strand twists about 90 degrees with exposed backbone polar atoms before entering into the protein core. While quite exposed, these target edge strands have distinctive structures that we hypothesized could allow for specific beta-strand complementarity.

Beta-strand interface conditioned RFdiffusion resulted in a 5-fold improved in silico success rate for these targets compared to hotspot directed RFdiffusion which, even without conditioning, generated beta-strand pairing binder designs with up to 20% of outputs (beta-strand pairing designs were generated <5% of the time for the less exposed edge strand containing targets). Binders for ɑ-CTX and NRP1 (1.9 nM and 84 nM respectively) were obtained by testing the 96 designs based solely on in silico metrics. The best binder to FCRL5, with a 82 nM Kd, was obtained from a yeast display library of 4846 designs. The FCLR5 and NRP1 target beta-strands both had similar conformational twists, but FCLR5bp and NRP1bp utilize quite distinct antiparallel and parallel hydrogen bonding binding modes, respectively (Figure 2a, right). Similar to FCLR5bp, ɑ-CTXbp forms a mostly canonical antiparallel strand pair with a slight irregularity as the binder beta-strand adapts to complement a small target strand bulge.

### Type III Receptor Tyrosine Kinases: KIT and PDGFRɑ

The type III RTK family receptors KIT and PDGFRɑ play roles in angiogenesis, tissue regeneration, and aberrant cancer signaling. Both receptor ectodomains are comprised of five Ig-like folds, with native ligands – stem cell factor and platelet derived growth factor – activating cellular signaling pathways by binding Ig domain 2 of KIT and PDGFRɑ, respectively, to induce receptor dimerization and intracellular cross-phosphorylation. Binders designed to occupy the ligand-binding pocket could act as antagonists to prevent aberrant signaling, and when oligomerized^50^, high-affinity binders could function as strong signaling agonists for tissue repair therapeutics. As such, we targeted the ligand binding sites on domain 2 of each receptor. As in the case of ALK-2 and ALK-3, previous attempts to de novo design binders against KIT had failed.

For KIT, a 75 nM binder was identified from a yeast surface library screening of 1230 designs. KITbp, with a target binding affinity of 64.7 nM, was designed to bind domain 3 strand on KIT which is part of the stem cell factor ligand binding site. The KITbp interface features an extensive canonical antiparallel hydrogen bond network, with 16 total residues forming 8 consecutive hydrogen bonds (Figure 2a, left). For PDGFRɑ, two binders were identified – PDGFRɑbp-7LBF (*K*_d_ = 137 pM) was designed to bind the a cryo-EM structure (PDB accession code: 7LBF), while PDGFRɑbp-AF2model (*K*_d_, 4 nM) was designed to bind the AlphaFold model. They were enriched from 3780 and 391 member design libraries via yeast display, without any further experimental sequence optimization. Despite significant disagreement between the cryo-EM structure and the AlphaFold model regarding the conformation of the target domain 2 Ig fold, design against both conformations yielded high-affinity binders. The binders may selectively induce the cognate PDGFRɑ conformation upon binding, as their design model conformations are not cross-compatible (Extended data figure 6). PDGFRɑbp-7LBF forms a complex h-bond network in which the binder strand forms both parallel and antiparallel interactions with two different target strands of the target domain 2 Ig fold. This precisely complementary strand pairing h-bond network highlights the power of beta-pairing conditioned RFdiffusion to design complex beta-strand architectures to perfectly complement idiosyncratic target topologies.

### Stability and solubility of designed binders with exposed edge beta-strand interfaces

No obvious trends were observed between the size of the edge beta-strand interface and aggregation propensity of the binders. All binders could be purified at high yields, and analysis of size exclusion chromatography curves reflect elution patterns consistent with monomeric binders being the most prominent purified species (Extended data figure 4).

Circular dichroism thermal melts (Figure 2b) obtained for each binder indicate that the binder folds remain intact at high temperatures, even those with significant edge beta-strand content. ALK-3bp was stable and monomeric up to 95 °C during a circular dichroism thermal melt experiment despite having four exposed edge beta-strands. The binder with the second most beta-strand content, ALK-2bp, was also thermostable with a measured TM of 81.2 °C. Binders with majority ɑ-helical support for their beta-strand interfaces (KITbp, ɑ-CTXbp, and PDGFRɑbp-AF2model) did not seem to be intrinsically more thermostable with measured TMs of 63.2 °C, 95 °C, and 95 °C respectively. The binders with ɑ/beta secondary structure content (2-3 beta-strand sheets with 2 buttressing ɑ-helices; FCRL5bp, NRP1bp, and PDGFRɑbp-7LBF) had TMs of >95 °C, 77.2 °C, and 71.1 °C, respectively. The excellent solution behavior of the designs despite having edge beta-strands clearly available for intermolecular interactions suggests that the same idiosyncratic features that enable high affinity and specificity (see below) target binding disfavor self-self interactions^9^.

### Beta-strand pairing binder interactions are target specific in all-by-all SPR assays

We next investigated whether the identified binders were specific for their designed target and did not form off target high affinity beta-strand pairing interactions. To test this, we performed an all-by-all SPR experiment where each binder was tested for binding affinity against each of the receptor targets in the test set. At 1.5 μM and 200nM concentrations, all of the binders showed strong SPR response for their intended target receptor compared to off-target receptors (Figure 3a). There was no evidence for off-target binding of the ALK-2, ALK-3, KIT, and PDGFRɑ binding proteins to related family members (e.g., ALK-2bp did not bind strongly to ALK-3). This high specificity may arise because the binders for each structurally similar target pair (i.e., ALK-2 and ALK-3, KIT and PDGFRɑ, FCRL5 and NRP1), were designed with different beta-strand pairing hydrogen bonding arrays, and each binder makes additional non-strand-pairing contacts with the intended target. With the exception of ALK-3bp which forms exclusively beta-strand contacts with ALK-3, all designed binders form additional ɑ-helical contacts that are complementary to neighboring hydrophobic surfaces adjacent to the target strands.

### Structural characterization of a KITbp:KIT complex confirms design model accuracy

To assess the accuracy of our design method, we solved the structure of KITbp in complex with domains 1-3 of KIT at 2.8 Å resolution (Figure 4a, Extended data table 1). The crystal structure has near exact agreement with the computational design model with 1.9 Å all-atom RMSD between the design model and the structure. Over the designed binder alone, the RMSD over all atoms between the crystal and design is 2.0 Å, and over the backbone, 1.0 Å RMSD (Figure 4b). The design model and crystal structure align with atomic level accuracy over nearly all interfacial side chain residue atoms. The KITbp binding site on KIT overlaps with that of the KIT native ligand stem cell factor (SCF), and consistent with this, saturating concentrations of SCF (20μM) reduced KITbp binding by >99% in a yeast surface binding assay.

There are 21 sidechain-sidechain and backbone-backbone hydrogen bonds between KITbp and KIT (calculated by HBPLUS^61^), many more than in previous designed binders and in most native protein-protein interfaces with similar (∼1500 Å^2^) interface sizes^62^ (Figure 4c). Of the 21 observed hydrogen bonds, 16 were predicted in our design model. At the center of the interface, a network of four binder and KIT tyrosine residues (Y38, Y39, Y259, Y269) form a highly complementary interface between the binder interfacial helix and core-boundary of the KIT domain 2 Ig fold. Lining the perimeter of the interface are numerous polar interactions, including the designed beta-pairing interface as well as several polar side-chain interactions.

KITbp contains an unpaired beta-strand that pairs with the target, with three buttressing ɑ-helices that tether the beta-strand in place. Foldseek^63^ alignment of KITbp against the PDB did not identify significant matches to known protein structures. Lone beta-strands are rare in nature and are found primarily in protein-protein complexes, where a loop in one partner forms a beta-strand conformation that extends the beta-sheet of the other partner. A similar induction of beta strand formation upon binding may occur with KITbp; Alphafold 2 prediction of the monomeric KITbp without KIT target places the beta-strand pairing residues in a coil conformation that better shields the binder’s hydrophobic core residues and hydrogen bonding atoms of the beta-strand interface (Extended data figure 7); such KIT-dependent conformational switching may contribute to the observed binding specificity.

### Biological functionality of designed binders

We next sought to assess the biological functionality of the designs. FCRL5 is internalized and transits through the endocytic degradation pathway upon binding to antibodies^62^, and we investigated whether FCRL5bp could be similarly internalized. We incubated cells expressing FCRL5 with FCRL5bp tagged with pHrodo DeepRed, a pH sensitive (∼5 pK_a_) dye that emits 655 nm fluorescence at late endocytic vesicle pH, and observed robust binder internalization that correlated with FCRL5 expression levels (Extended data figure 8a), reaching a steady state at in this expression system. FCRL5bp could thus be useful as an Endotag^52^ for targeted protein degradation, as recently demonstrated for other designed proteins. PDGFRɑbp-7LBF was designed to sterically occlude the binding site of the native ligand PDGF-AA, and indeed PDGFRɑbp-7LBF blocked activation of signaling by PDGF-AA through the Akt and Erk pathways with nanomolar IC50s (Figure 5). In a companion study, ɑ-CTXbp was found to potently neutralize ɑ-CTX, protecting mice from a lethal neurotoxin challenge^64^.

## Discussion

We show that beta-strand interface conditioned RFdiffusion outperforms previous methods for the design of binders against edge beta-strand containing targets. The binders are thermostable and target-specific, and the crystal structure of the KITbp:KIT complex shows the method has high structural accuracy. The higher success rate of binder design against largely polar targets containing beta-sheets using the beta-pairing targeted approach than with primarily ɑ-helical designs likely reflects the difficulty for the latter of complementing the many polar NH and CO groups on edge beta-strands with designed sidechain-backbone hydrogen bonds–using geometrically matched beta-strands this polar interface can be achieved in less convoluted fashion. The very high binding specificity and the lack of self association of our designs despite the availability of edge beta-strands for intermolecular interactions support the conjecture of the Richardsons^9^ that irregular beta-strand geometries suppress off target pairings. Our beta-strand targeted RFdiffusion approach should enable facile binder design against many previously challenging protein surfaces, including the many Ig domains and other beta-sheet containing domains frequent in the extracellular domains of cell surface receptors; as many of these are current therapeutic targets, the method could contribute to new therapeutic interventions.

## Methods

### Computational protein design workflow

Target structures used as inputs for binder design were obtained from the Protein Data Bank for KIT (PDB:2E9W)^65^, NRP1 (PDB:2QQI)^66^, and PDGFRɑ (PDB:7LBF)^67^. Publicly available alphafold models were used for the design of binders against ALK-2(Uniprot:Q04771), ALK-3 (Uniprot:P36894), FCRL5 (Uniprot:Q96RD9), and PDGFRɑ (Uniprot:P16234). Edge beta-strand target residues were selected to condition RFdiffusion towards generating beta-strand interfaces with these edge strands. ProteinMPNN sequences were generated for the output backbones and subject to in silico screening based on AlphaFold2 initial guess (pAE interaction and binder pLDDT)^6^, AlphaFold2 monomer pLDDT^59^, Rosetta ΔΔG^68^, radius of gyration, and spatial aggregation propensity^69^. The cutoff values varied for each target protein. Generally, RFdiffusion backbones were generated at the 10^4^ scale, and multiple partially diffused perturbations and ProteinMPNN sequences for the initial RFdiffusion backbones were screened in silico at the 10^5^ scale. To ensure our binder design libraries sampled diverse sequences and structures, we clustered designs by sequence similarity or RMSD and selected a subset of designs such that each cluster was equally represented in the final library for experimental characterization.

### DNA library preparation

For ALK-2, ALK-3, KIT, FCRL5, and PDGFRɑ, 10^4^ scale DNA libraries were generated by using DNAworks2.0 reverse translating designed amino acid sequences that optimally reflected the Saccharomyces cerevisiae codon frequency table. Additional 5’ and 3’ adapters were included to enable PCR amplification of libraries by single sets of primers. All libraries were amplified using Kapa HiFi polymerase (Kapa Biosystems) with a qPCR machine (Bio-Rad, CFX96). In detail, the libraries were first amplified in a 25 μl reaction, and the PCR reaction was terminated when the reaction reached half maximum yield to avoid overamplification. The PCR product was loaded onto a DNA agarose gel. The band with the expected size was cut out, and DNA fragments were extracted using QIAquick kits (Qiagen). Then, the DNA product was re-amplified as before to generate enough DNA for yeast transformation. The final PCR product was cleaned up with a QIAquick Clean up kit (Qiagen). For the yeast transformation step, 2–3 µg of linearized modified pETcon vector (pETcon3) and 6 µg of insert were transformed into the EBY100 yeast strain using a previously described protocol.

DNA libraries for deep sequencing were prepared using the same PCR protocol, except the first step started from yeast plasmid prepared from 5 × 10^7^ to 1 × 10^8^ cells by Zymoprep (Zymo Research). Illumina adapters and 6-bp pool-specific barcodes were added in the second qPCR step. Gel extraction was used to obtain the final DNA product for sequencing. All the different sorting pools were sequenced using Illumina NextSeq sequencing.

### Target protein preparation

Biotinylated target protein was commercially available for KIT (Acro Biosystems, CD7H5255), FCRL5 (Acro Biosystems, FC5-H82E3), NRP1 (Acro Biosystems, NR1-H82E3), α-CTX (Latoxan, L8114), and PDGFRɑ (Sino Biological, 10556-H27H-B). For ALK-2, ALK-3 biotinylated receptor was expressed as avi-tagged ectodomain constructs in *E. coli*, purified, and biotinylated as described by Tao Huang and Andrew P. Hinck^70^.

### Yeast Display

Saccharomyces cerevisiae EBY100 strain cultures were grown in C-Trp-Ura medium supplemented with 2% (w/v) glucose. For induction of expression, yeast cells were centrifuged at 4,500 g for 5 min and resuspended in SGCAA medium supplemented with 0.2% (w/v) glucose at the cell density of 1 × 10^7^ cells per ml and induced at 30 °C for 16–22 h. Cells were washed with PBSF (PBS with 1% (w/v) BSA) and labeled with biotinylated targets using two labelling methods: with-avidity and without-avidity labeling. For the with-avidity method, the cells were incubated with biotinylated target, together with anti-c-Myc fluorescein isothiocyanate (FITC, Miltenyi Biotech) and streptavidin–phycoerythrin (SAPE, ThermoFisher). The concentration of SAPE in the with-avidity method was used at one-quarter of the concentration of the biotinylated targets. For the without-avidity method, the cells were first incubated with biotinylated targets, washed and secondarily labeled with SAPE and FITC. All the original libraries of de novo designs were sorted using the with-avidity method for the first few rounds of screening to exclude weak binder candidates, followed by several without-avidity sorts with different concentrations of biotinylated targets.

### Protein expression and purification

Protein binder designs were ordered as synthetic genes (eBlocks, Integrated DNA Technologies) and cloned via BsaI overhangs into the target cloning vector, LM0627^71^ for Golden Gate assembly. Golden Gate reaction mixtures were transformed into a chemically competent expression strain (BL21(DE3)), and overnight outgrowth cultures were used to seed 500mL protein expression cultures in auto-induction media (autoclaved TBII media supplemented with Kanamycin, 2mM MgSO4, 1X 5052). The following day (20-24 hrs later), cells were harvested and lysed by sonication (QSonica Q500 Sonicator), and clarified lysates were purified by immobilized metal affinity chromatography using Ni-NTA agarose resin (Qiagen). Samples were eluted in a Tris elution buffer containing 300mM imidazole, sterile filtered with 0.22μm Polyvinylidene Fluoride syringe filter prior to size exclusion chromatography. Protein designs were then screened via SEC using an AKTA FPLC outfitted with an autosampler capable of running samples from a 96-well source plate. The protein binders were run on a Superdex75 Increase 5/150 GL column (Cytiva 29148722).

### Circular Dichroism

Far-ultraviolet circular dichroism measurements were carried out with a JASCO-1500 instrument equipped with a temperature-controlled multi-cell holder. Wavelength scans were measured from 260 to 190 nm at 25 and 95 °C. Temperature melts monitored the dichroism signal at 217 nm in steps of 2 °C min–1 with 30 s of equilibration time. Wavelength scans and temperature melts were performed using 0.3 mg ml–1 protein in PBS buffer (20 mM NaPO4, 150 mM NaCl, pH 7.4) with a 1 mm path-length cuvette. Melting temperatures were determined by fitting the data with a sigmoid curve equation. For designs retained more than half of the mean residue ellipticity values, Tm values are reported as greater than 95 °C. Tm values of the other designs were determined as the inflection point of the fitted function.

### Surface plasmon resonance for determination of binding kinetics

Binding kinetics were analyzed via Surface Plasmon Resonance (SPR) on a Biacore 8K (Cytiva). Binding to receptors was measured by capturing biotinylated receptor ectodomain on a Streptadvidin coated chip using Biotin CAPture Kit (Cytiva #28920234). by injecting 0.125 µg/mL receptor at a flow rate of 10 µL/min in HBS-EP+ (0.01 M HEPES pH 7.4, 0.15 M NaCl, 3 mM EDTA, 0.005% v/v Surfactant P20, Cytiva #BR100669). Analytes were diluted in HBS-EP+ and injected at a flow rate of 30 µL/min to monitor association. HBS-EP+ was used as a running buffer during dissociation at a flow of 30 µL/min. Full kinetics by running single cycle kinetics, injecting increasing concentrations of ligand or parallel kinetics using different concentrations in the eight channels. Association/dissociation times and concentration ranges were varied to suit the respective analytes. Binding kinetics were determined by global fitting of curves to k_on_ and k_off_ assuming a 1:1 Langmuir interaction, using the Cytiva evaluation software.

### Surface plasmon resonance for measuring binder-target specificity

All by all binder-target interactions were measured according to SPR protocol described above. Identical binder titres and receptor loading protocols were used to maintain as much signal consistency as possible, with binder titre values ranging in seven fold dilutions from 10 μM to 600pM to capture nonspecific binding across a large concentration spectrum. Non-specific binding responses were evaluated by averaging response values across the association phase for a given titre, normalizing responses such that all values were positive, and taking each response as a fraction of the maximum observed response (for each binder the maximum response was observed against the target protein for which it was designed).

### Recombinant production of KIT_D1-D3_ for X-ray crystallography

KIT123, i.e., extracellular domains D1–3 of the tyrosine kinase receptor KIT, was recombinantly produced via transient expression of suspension-adapted HEK 293 cells. Cells were grown and maintained in a 1:1 mixture of the Freestyle (Gibco) and the Ex-Cell (Merck) medium. Before the transfection, cells at the density of 1.5×10^6^ cells.mL^-1^ were centrifuged at 250×g for 6 min and resuspended in the pre-warmed FreeStyle medium only to reach the density of 3.0×10^6^ cells.mL^-1^ and incubated at 37 °C, 130 rpm, 70 % humidity, and 8.0 % CO_2_ for 15 min. The cells were subsequently added 450 µg of the plasmid DNA carrying the target construct per 100 mL of the medium. After 5 min, the cells were added 900 µg of linear polyethylenimine 25 kDa (Polysciences) and 3 µmol of kifunensine per 100 mL of the medium and continued incubation. After additional 5 h, an equal volume of the Ex-Cell medium to the FreeStyle medium was added to the cultures to return back to the density of ∼1.5×10^6^ cells.mL^-1^. 24 hours post-transfection, the cells were added D-glucose and valproic acid to the resulting concentration of 55 mM and 3.5 mM, respectively. 96 hours post-transfection, the cells were harvested by centrifugation at 500×g and 4 °C for 10 min and the conditioned medium (supernatant), carrying the recombinantly produced protein of interest, was collected, added 10,000 U of Endo Hf (NEB) per 100 mL of the medium, and incubated for 4 h at room temperature to remove heterogenous N-linked glycans and facilitate the subsequent crystallization attempts. After filtering the incubated medium through a 0.22 µm filter, the clarified sample was loaded to a 1 mL Ni-NTA HisTrap HP column (Cytiva) equilibrated with HEPES-buffered saline (HBS; 20 mM HEPES, 150 mM NaCl, pH 7.4). The column was washed with 10 mM imidazole in HBS and the His-tagged protein of interest was eluted using 150 mM imidazole in HBS. The eluted protein fraction was subsequently loaded to a Superdex 75 Increase 10/300 GL column (Cytiva) to simultaneously remove aggregates and remaining impurities and to exchange buffer to HBS with no imidazole. The fractions corresponding to the protein of interest were pooled together, their purity was analyzed by SDS-PAGE (Bio-Rad), and the concentration was determined using the NanoDrop Spectrophotometer (Thermo Fisher Scientific).

### Crystal structure of KIT123 in complex with KITmb

The KIT123–KITbp complex was formed by adding a 3-fold molar excess of the purified KITbp to the recombinantly produced, EndoH-treated KIT123 receptor (domains D1–3 of the ectodomain). The complex was isolated using size-exclusion chromatography equipped with a Superdex 75 Increase 10/300 GL column (Cytiva) equilibrated with HEPES-buffered saline (HBS; 20 mM HEPES, 150 mM NaCl, pH 7.4). Fractions corresponding to the KIT123–KITbp complex were pooled and concentrated by centrifugal ultrafiltration to the concentration of 6.1 mg.ml^-1^. Sparse-matrix crystallization screens were carried out in 96-well 3-drop plates (Molecular Dimensions) using the BCS-Screen (Molecular Dimensions) at 293 K and the sitting-drop method. The vapour-diffusion geometry was used to set up sitting drops consisting of 100 nL of a protein solution and 100 nL of each reservoir solution using a Mosquito nanolitre crystallization robot (SPT Labtech). The protein complex crystallized in the condition G11 (0.2 M sodium/potassium phosphate pH 7.5, 0.1 M HEPES pH 7.5, 22.5 % PEG Smear Medium, 10 % glycerol). Crystals were cryo-protected with mother liquor supplemented with 25% v/v glycerol and subsequently flash-cooled by direct plunging into liquid nitrogen. X-ray diffraction data of protein crystals were collected at the P13 beamline (PETRA III, EMBL Hamburg). Obtained data were processed using XDS^72^ and severe data pathologies, including strong anisotropy and translational noncrystallographic symmetry, were revealed, yielding similar characteristics as reported previously^73^. Based on these findings, the data were elliptically-truncated and corrected using the STARANISO^74^ server and accordingly treated during the following steps. Initial phases were determined by maximum-likelihood molecular replacement in Phaser^75^ using the domains D1–3 part of the KIT-SCF structure (PDB ID: 2E9W)^65^ as a search model. Model (re)building was performed in Coot^76^, and coordinate and ADP refinement was performed in PHENIX^77^.

Model and map validation tools in Coot, the PHENIX suite, and the PDB_REDO server^78^ were used to validate the quality of crystallographic models. Atomic coordinates and structure factors of the protein-protein complex were deposited in the Protein Data Bank^79^ under the PDB code 9H71.

### PDGFRɑ antagonism assay

Heparan-deficient Chinese hamster ovary cells stably overexpressing PDGFRα (CHO-PDGFRα) were grown to 70-90% confluency in CHO growth medium (Kaighn’s Modification of Ham’s F12 (F12K) medium (ATCC# 30-2004) + 10% Fetal Bovine Serum (FBS) (Biowest, #S1620) + 4% Penicillin-Streptomycin (P/S) (Gibco, #15140122) with 10 μg/mL puromycin (Gibco, #A11138-03). The cells were starved for 4 hours in serum free F12K media and treated with synthetic and/or native ligand for 15 minutes at 37 degrees celsius. Cells were subsequently washed with PBS and lysed with buffer containing 20 mM Tris–HCl (Sigma-Aldrich, 1185-53-1) (pH 7.5), 150 mM NaCl, 15% glycerol (Sigma-Aldrich, G5516), 1% triton (Sigma-Aldrich, 9002-93-1), 3% SDS (Sigma-Aldrich, 151-21-3), 25 mM β-glycerophosphate (Sigma-Aldrich, 50020-100G), 50 mM NaF (Sigma-Aldrich, 7681-49-4), 10 mM sodium pyrophosphate (Sigma-Aldrich, 13472-36-1), 0.5% orthovanadate (Sigma-Aldrich, 13721-39-6), 1% PMSF (Roche Life Sciences, 329-98-6), 25 U benzonase nuclease (EMD, 70664-10KUN), protease inhibitor cocktail (PierceTM Protease Inhibitor Mini Tablets, Thermo Scientific, A32963), and phosphatase inhibitor cocktail 2 (Sigma-Aldrich, P5726). Lysates were collected, mixed with 4x Laemmli buffer (BioRad, #161-0747), and boiled at 95 Celsius for 10 minutes before 10 uL were loaded on to 4-10% SDS-PAGE gels and run for 30 min at 250 Volts.

### Western blotting

The PDGFRɑ antagonism assay was analyzed using two different western blot techniques. One repeat of each was analyzed via traditional techniques which is as follows: after separation, proteins were transferred onto a nitrocellulose membrane (12 minutes, semi-dry transfer) and blocked for one hour in 5% bovine serum albumin. Membranes were probed with the following primary antibodies: phospho-PDGFRα (Tyr762) (Cell Signaling Technology, #24188) 1:1000 dilution, Phospho-Akt (Ser473) (Cell Signaling Technology, #9271) 1:1000 dilution, Phospho-p44/42 MAPK (Erk1/2) (Thr202/Tyr204) (Cell Signaling Technology, #9101) 1:10,000 dilution, and either S6 (Cell Signaling Technology, #2117) at 1:1000 dilution or H3 (Cell Signaling Technology, #9715) at 1:5000 dilution as a loading control. After overnight incubation on a rocker, the membrane was probed with HRP-conjugated secondary antibody, washed 3 times, and imaged with a Bio-Rad ChemiDoc Imager. The blot image was quantified using ImageJ peak band intensity. In addition, samples were analyzed using a Biotechne Jess Automated Western Blot Machine (Biotechne, #004-650). Samples were diluted 1:3 before being prepared as per the kit instructions for 12-230 kDA separation modules (Biotechne, #SM-W001). Samples were probed using the same primary antibodies as above via a RePlex assay (Biotechne, #RP-001) that was run using the default settings for anti-rabbit chemiluminescence (Biotechne, #DM-001). Signal was quantified using area under the curve for each protein of interest. All data points from each assay were normalized to a housekeeping gene in the same lane and and then normalized to the 1 nM PDGF-AA condition within each experiment.

### Vectors and constructs/ Lentiviral generation and infection

Lentiviral particles of the pHAGE-PDGFRɑ plasmid (Addgene, 116769)^77^ were generated by transfecting 60-80% confluent HEK293FT cells, maintained in 10 mL HEK growth medium (Gibco™ DMEM, high glucose (Gibco, 11965092) + 10% FBS + 1% P/S) on a 100mm TC-treated culture dish, with 20 µg pHAGE-PDGFRɑ, 15 µg psPAX2 (Addgene, 12260), and 5 µg pMD2.g (Addgene, 12259) combined with 1.8 mL Opti-MEM medium (Gibco, 31985070) and 60 µg linear polyethylenimine. Transfected HEK293FT cells were replenished with fresh HEK growth medium 24 hours after the transfection. The supernatant containing lentivirus was collected 48 and 72 hours post-transfection. The collected supernatant was filtered through a 0.45 µM PES syringe filter. CHO-PDGFRα cells were generated by infecting 60-80% confluent heparan-deficient Chinese Hamster Ovary cells (CHO) cells (pgsD-677 cells) (# CRL-2244) with the filtered lentivirus-containing supernatant. Infected CHO cells were supplemented with fresh CHO growth medium 24 hours after the lentiviral infection. 48 hours after the infection, CHO cells were selected in the CHO selection medium (CHO growth medium containing 10 µg/mL puromycin). CHO selection medium was replaced every 48 hours for 7 days. CHO-PDGFRα cells post-selection were maintained in the selection medium.

### FCRL5 cell line preparation

Doxycycline-inducible expression of Flag-FCRL5-P2A-T2A-EGFP was generated by first seeding 1 x 10^6^ WT HeLa cells in 10 cm dish (Genesee 25-202) in Dulbecco’s Modified Eagle Medium (DMEM, Gibco 11995073) supplemented with 10% heat-inactivated fetal bovine serum (HI FBS, Gibco A5256801) and 1% penicillin-streptomycin (PS, Gibco 15140122). The next day, cells were transfected using TransIT-LT1 Transfection reagent (Mirus Bio MIR2300) according to manufacturer protocol with 2:1 donor:sleeping beauty transposase plasmids (Addgene Plasmid #34879)^80^. After 72 h, cells were covered with selection media comprised of DMEM + 10 % HI FBS + 1% PS and 2 μg/mL puromycin (Invivogen 58-58-2). Selection media was replaced every 48 h until control WT cells were completely dead. Cells were maintained in selection media.

### FCRL5bp internalization assay

Flag-FCRL5 HeLa cells were counted and seeded at 50,000 per well in a 24-well plate (Genesee 25-107) in DMEM + 10% HI FBS + 1% PS. The next day, cells were treated with either 0, 100, or 1,000 ng/mL doxycycline (Fisher Scientific BP26531), and incubated for 48 h at 37 C 5% CO_2_. Cells were then lifted, counted using TrypanBlue, and seeded at 12,500 cells per well in a 96-well plate (Corning 3595) in phenol red-free DMEM (Gibco 31053028) supplemented with 10 % HI FBS and 0, 100, or 1,000 ng/mL. Cells were incubated at 37 C 5% CO_2_ for 8 h. Then, AviTagged-FCRL5bp was diluted in phenol red-free DMEM+10% HI FBS and respective doxycycline concentrations. Then, FCRL5bp was complexed 1:1 with TFP ester-pHrodoDeepRed (Invitrogen P35358) labeled NeutrAvidin (Thermo Scientific 31000) for 15 min at 37 C covered from light. Media was then replaced with respective treatments, and cells were monitored with a live-cell imaging incubator (Sartorius Incucyte S3) and internalization was quantified as the overlap of red and green fluorescent area divided by the total phase area in a well.

## Acknowledgements

This research was supported by The Audacious Project at the Institute for Protein Design (I.S., D.L. S.V.T., J.N.S., X.W., D.B.), NSF-GRFP program (S.A.R.), Howard Hughes Medical Institute (D.B., I.G., B.C., M.G.S, N.R., D.V., M.G., J.N.S.), The Nordstrom Barrier Institute for Protein Design Directors Fund (B.H.), The Open Philanthropy Project Improving Protein Design Fund (S.V.T. and B.C.), and AMGEN Donation to the Institute for Protein Design (X.W.). The project or effort depicted was or is sponsored by the Department of the Defense, Defense Threat Reduction Agency grant HDTRA1-21-1-0007 (M.G., X.W.). S.N.S. is supported by the Flanders Institute for Biotechnology (VIB), Belgium (grant no. C0101), Research Foundation Flanders (FWO), Belgium (grant nr. S000722N), and an FWO EOS research grant (grant nr. G0H1222N). M.T. has received funding from the European Union’s Horizon Europe research and innovation programme under the Marie Skłodowska-Curie grant agreement No. 101155448 (HeartRepairKIT). T.P.J. acknowledges support from the Alliance programme under the EuroTech Universities agreement. M.B.V. has received funding from the European Union’s Horizon 2020 research and innovation programme under the MarieSkłodowska-Curie grant agreement No 899987. Biotinylated ALK-2 and ALK-3 were kindly provided by Andrew Hinck. We thank the staff of the beamline P13 (PETRA III, EMBL Hamburg) for technical support and beamtime allocation. We also acknowledge Preetham Venkatesh and Fiona Wang for support in testing the design methodology described in this study, Xinting Li for help with mass spectrometry analysis of proteins, Rohith Krishna for logistics support, and Ashish Phal for identifying targets of interest.

