## Extended Data for "Improved protein binder design using beta-pairing targeted RFdiffusion"

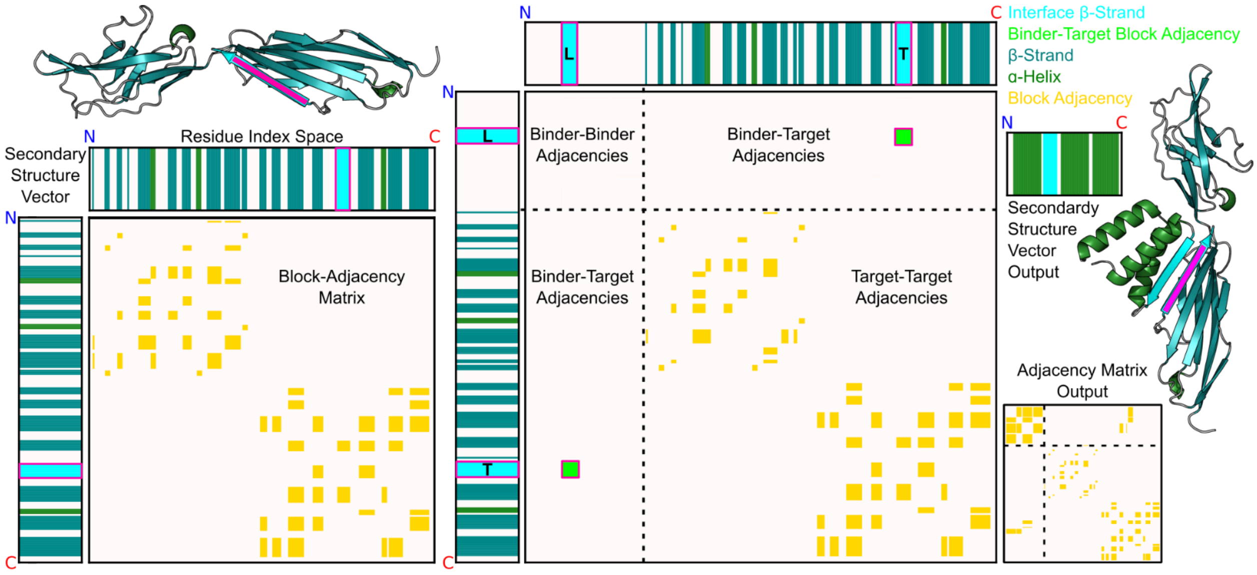


**Extended data Figure 1.** Design of beta-strand pairing scaffolds with RFdiffusion. To leverage the diverse generative potential of RFdiffusion while encouraging beta-strand interfaces in design outputs, we implemented an interface conditioning algorithm that generate SS/ADJ conditioning tensors given simple user inputs. The model understands fold conditioning in the form of tensors that label the secondary structures (blue) of each residue (a, top and left), and the adjacencies of these secondary structure blocks (a, yellow center). User specified parameters specify information about the binder-interfacial secondary structure block (a beta-strand in this case), the length of this block (b, cyan block in binder tensor L), and the target site residues to which the binder block is adjacent (b, cyan block in target tensor T). From these predefined parameters, the algorithm randomly samples the location of the binder-interfacial secondary structure block in residue index space, while maintaining a defined adjacency (green) to the specified target site residues. This user-defined conditioning tensor guides diffusion outputs towards a beta-strand pairing binder-target interface (c). Previously, RFdiffusion interface design calculations could be directed to specific residues specified as target “hotspots” to specify the intended target residues to bind to, this new inter-chain SS/ADJ conditioning enables user indication of “beta-strand hotspots" or “α-helical hotspots” during binder scaffold generation. Binder scaffold outputs conditioned on expanded binder-target SS/ADJ tensors enabled design of user-specified beta-strand interface types.

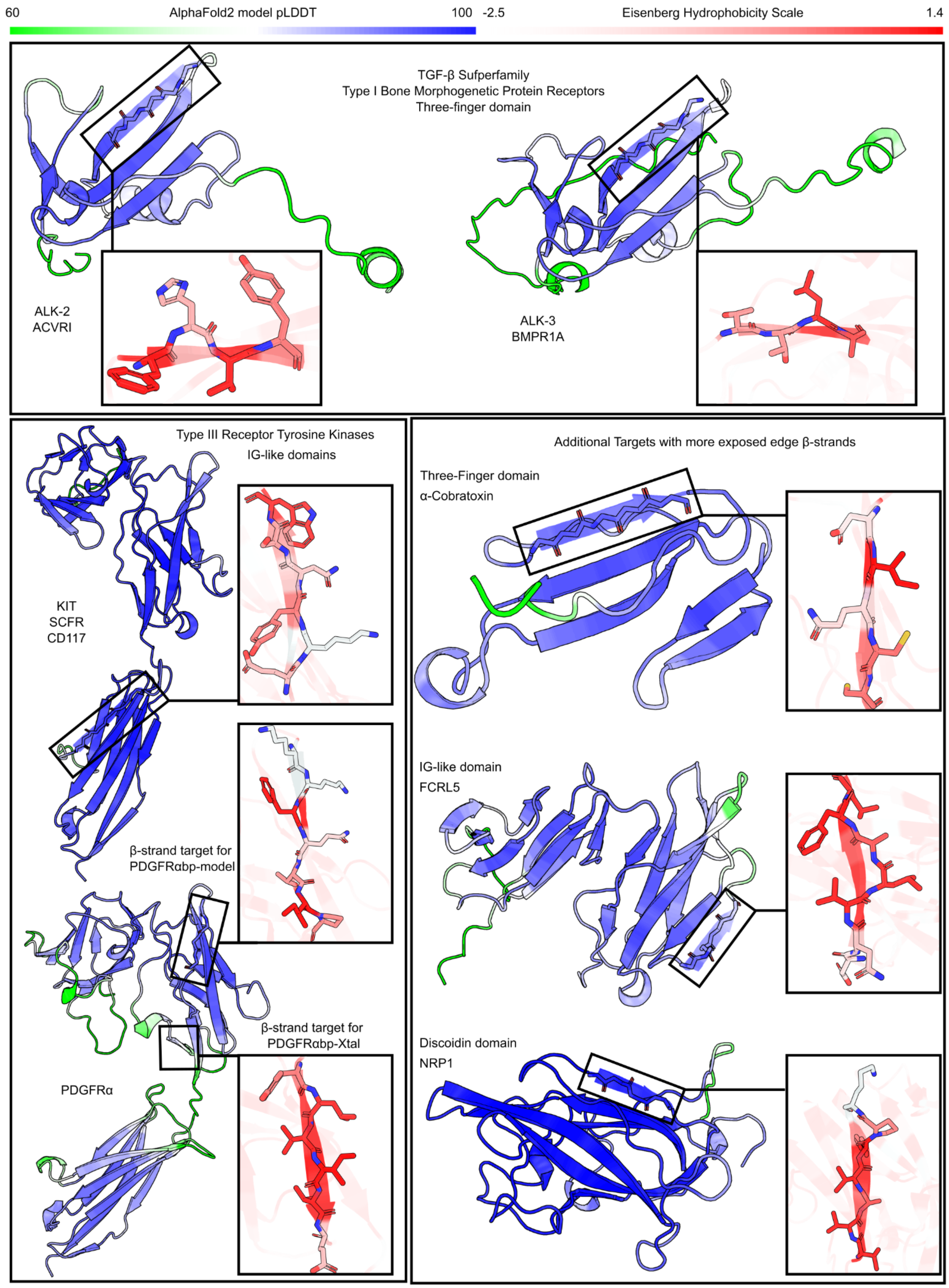
60
AlphaFold2 model pLDDT
100
-2.5
Eisenberg Hydrophobicity Scale
1.4
TGF-β Sufperfamily
Type I Bone Morphogenetic Protein Receptors
Three-finger domain
ALK-2
ACVRI
ALK-3
BMPR1A
Type III Receptor Tyrosine Kinases
IG-like domains
KIT
SCFR
CD117
β-strand target for
PDGFRαbp-model
β-strand target for
PDGFRαbp-Xtal
PDGFRα
Additional Targets with more exposed edge β-strands
Three-Finger domain
α-Cobratoxin
IG-like domain
FCRL5
Discoidin domain
NRP1

**Extended data Figure 2.** Edge beta-strands on target proteins as binding sites. In order to test interaction specificity of designed binders, target proteins with similar folds were selected. Three targets (ALK-2, ALK-3, and α-CTX) contain three-finger domains, with the N-terminal edge beta-strands targeted for ALK-2 and ALK-3 binders, and the C-terminal edge beta-strand targeted for α-Cobratoxin binders. Three other targets (KIT, PDGFRα, and FCRL5) contain immunoglobulin domains. Ig fold N-terminal strands were targeted for KIT and PDGFRα (7LBF) binders, and a non-terminal edge beta-strand was targeted on FCRL5. Additionally, we targeted the C-terminal beta-strand on PDGFRα (AF2-model) binders. Finally an edge beta-strand on a discoidin domain of NRP1 was targeted. Alphafold models are shown with per-residue pLDDT values ranging from 60 (green) to 100 (blue). High resolution experimentally determined structures (PDB IDs – ALK-2:7YRU, ALK-3:2H62, α-CTX:1CTX, KIT:2E9W, NRP1:2QQI, and PDGFRA:7LBF) of the targeted edge beta-strands are shown, with residues labeled by Eisenberg Hydrophobicity Scale (white are more polar and red are more hydrophobic). No experimental structure exists for FCRL5, and the Alphafold model edge beta-strand is shown here.

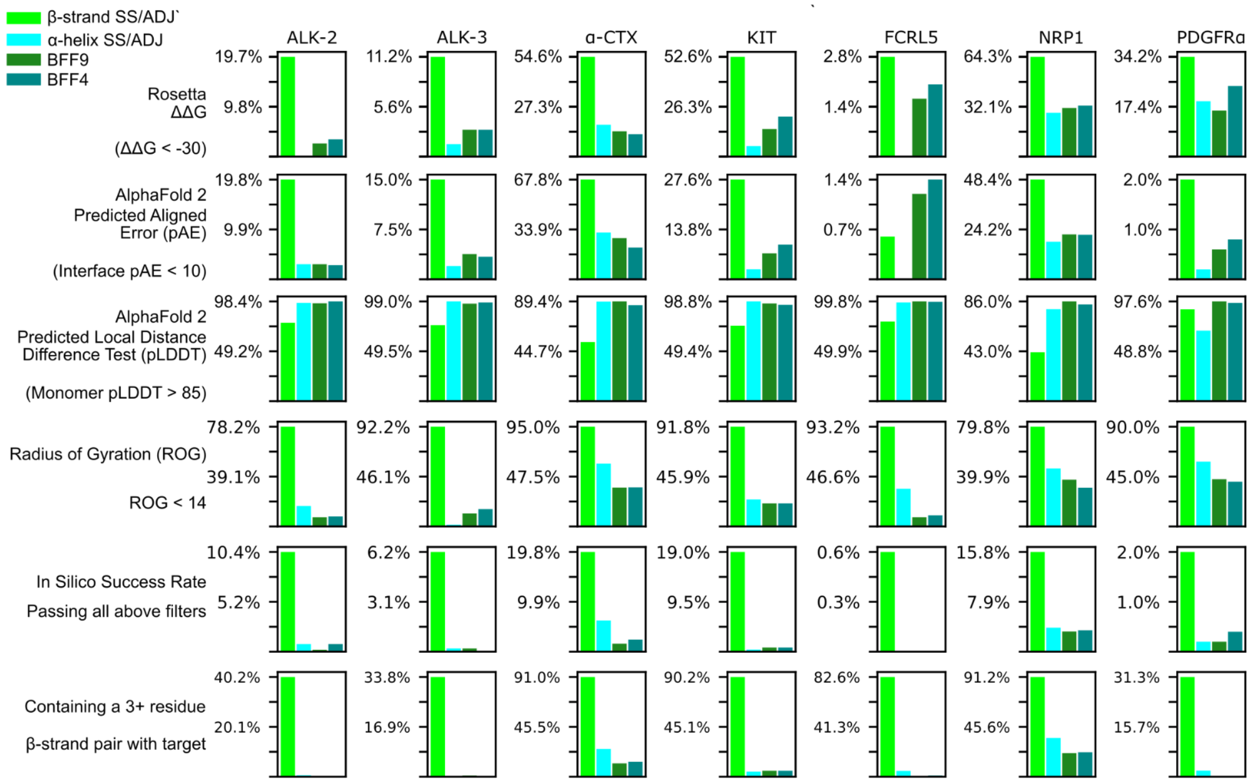


**Extended data figure 3.** Success rates from different RFdiffusion methods across several predictive in silico metrics. 40,000 total designs were generated, with equal numbers generated from different RFdiffusion scaffold generation methods across eight target structures. The methods used included hotspot-only conditioning by the BFF4 RFdiffusion checkpoint (teal, the default for hotspot conditioned binder design), hotspot-only conditioning by the BFF9 checkpoint (dark green, the checkpoint used for interface SS/ADJ conditioning), α-helix interface and hotspot conditioning (cyan), and beta-strand interface and hotspot conditioning (green).

Alphafold predictions yielded two in silico filtering scores: predicted error among interface residues (interface pAE) and overall confidence in the binder fold (pLDDT). Binders with interface pAE < 10 and binder pLDDT > 85 are considered successful designs with respect to these metrics. Rosetta ΔΔG represents the Rosetta predicted change in energy in the binder designs and targets upon binding. Lower ΔΔG is predictive of higher affinity interactions, with a ΔΔG value of -30 used as a standard of success here. Radius of gyration is a statistic of protein globularity that is predictive of a protein's fold stability. In nature, nearly all 65 amino acid proteins (the size of all designs in this in silico benchmarking set) have a radius of gyration < 14, mostly reflecting  known globular folds, such as 3+ α-helix bundles, mixed α/beta proteins, and beta-sandwich folds. Binders meeting each of these four criteria were considered in silico successes with a high likelihood of experimental success. Additionally, the percentage of outputs exhibiting binder-target beta-strand pairing are plotted, although this was not used as a filter for success.

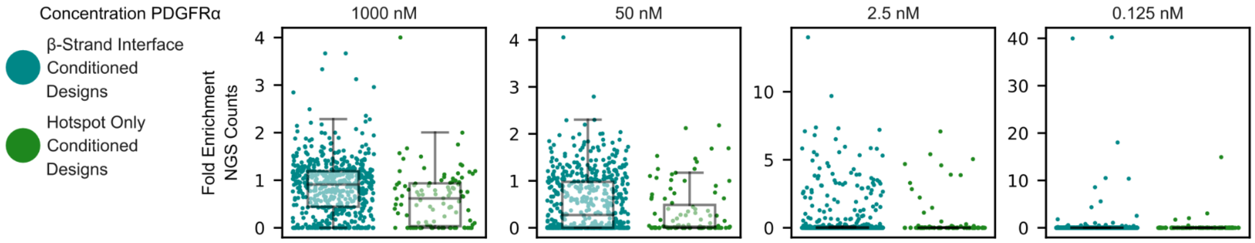


**Extended data Figure 4.** Beta-strand pairing PDGFRα binders are more enriched for target binding in yeast surface display. Two binder libraries, beta-strand interface conditioned (N=5616) and hotspot only conditioned (N=5717) designed against PDGFRα were expressed in vectors to display binders extracellularly and with a Myc-tag. Libraries were pooled, sorted for expression by binding to fluorescent Anti-Myc-FITC. The expression positive fraction was initially sorted for PDGFRα binding by incubating cells with biotinylated PDGFRα and Streptavidin phycoerythrin prior to sorting. This parent sort was repeated with titrated PDGFRα concentrations as labeled, and the fold enrichment of each binder in this sort was plotted relative to the NGS counts in the parent sort. Indeed the binders that ultimately showed the tightest binding when purified and tested by SPR were the two best performing yeast clones in this assay.

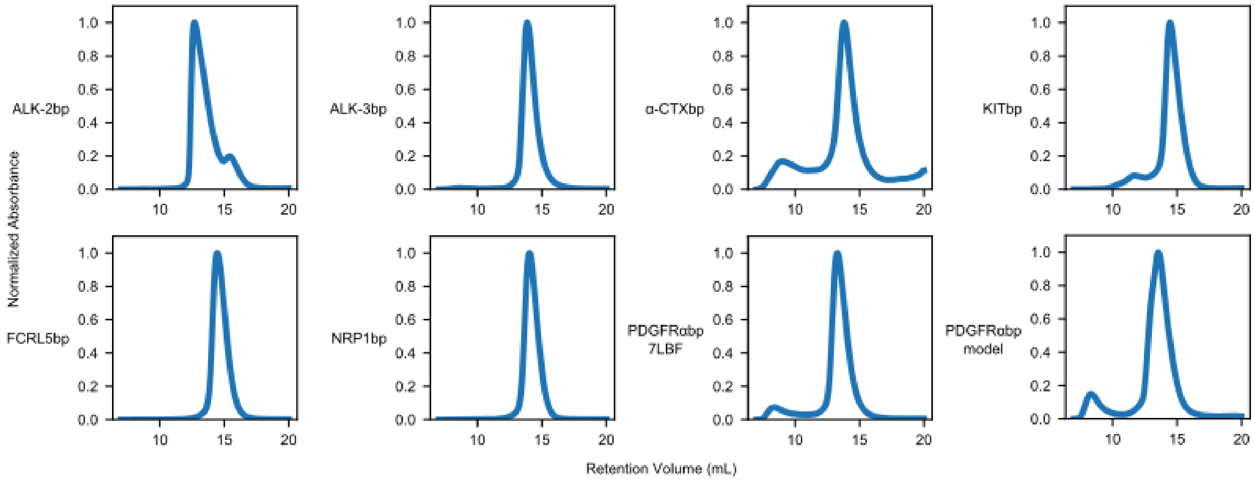


**Extended data Figure 5.** Purification of beta-strand pairing binders shows binder monodispersity. Each binder expressed in high yields (>1mg) from 500 mL *E. coli* cultures. Each trace is within the expected range for monomeric species. There is minimal evidence of significant aggregation, with aggregate peak areas (<10 mL retention volume) comprising less than 10% of the overall eluate for 7/8 designs.

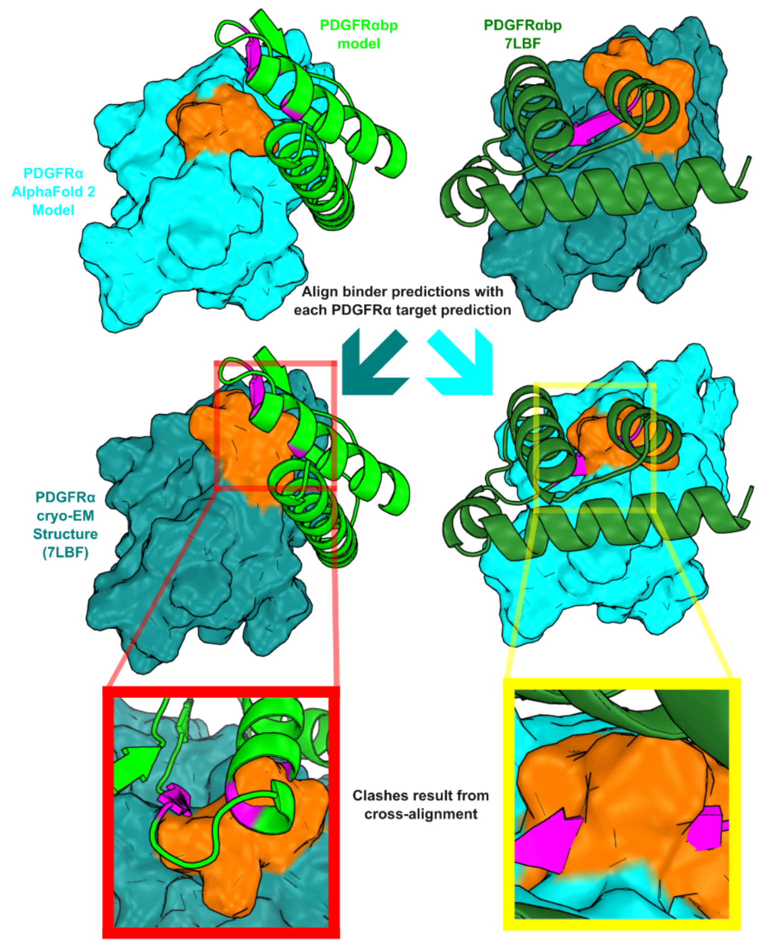


**Extended data figure 6.** The two PDGFRα binder design models are predicted to induce different PDGFRα conformations. Our PDGFRαbp-model binder (lime green) designed against the PDGFRα Alphafold 2 model (cyan) binds to target site residues (orange) in a conformation that significantly varies from the available cryo-EM structure (PDB ID 7LBF, teal). When the Alphafold 2 model binder (dark green) is aligned to the cryo-EM model, the binder interface clashes with the target site residues (red inset). Similarly, aligning the PDGFRAbp-7LBF binder (dark green) designed against the cryo-EM structure to the Alphafold 2 model results in a target site clash. Structure agnostic (i.e., only sequences are provided) Alphafold prediction of each binder in complex with PDGFRA reflects the conformational variability, with the Alphafold model conformation favored in PDGFRAbp-model predictions while the cryo-EM conformation is favored when predicted with PDGFRAbp-7LBF.

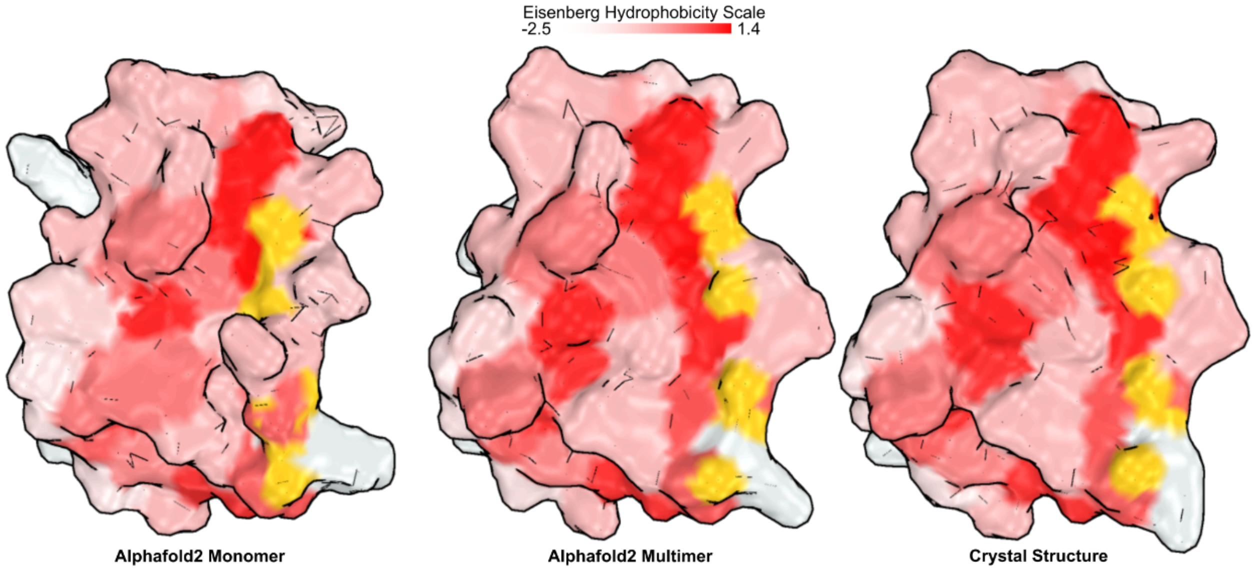


**Extended data figure 7.** Surface hydrophobicity representations of the KITbp interface for the monomeric (top) KIT-bound (middle) Alphafold2 predictions of KITbp. The crystal structure representation (bottom) and the Alphafold2 multimer prediction display more of KITbp hydrophobic residues than the Alphafold 2 monomer prediction. The hydrogen bonding atoms (gold) are observed in a beta-strand conformation amenable to target beta-strand pairing in the KIT-bound conformations, while these same atoms are occluded in the Alphafold 2 monomer prediction.

| **Crystal data** | |
|---|---|
| Space group | $P2_12_12_1$ |
| Unit cell parameters *a*, *b*, *c* [Å] | 41.273, 78.643, 291.008 |
| Unit cell parameters *α*, *β*, *γ* [°] | 90, 90, 90 |
| **Data collection** | |
| Wavelength [Å] | 0.9762 |
| Resolution range [Å] | 69.18–2.80 (2.90–2.80) |
| Total reflections | 250 107 (37 292) |
| Unique reflections | 24 388 (3 846) |
| Multiplicity | 10.3 (9.7) |
| Completeness [%] | 99.8 (99.4) |

| | |
|---|---|
| Mean I/σ [I] | 5.24 (0.69) |
| Wilson B-factor | 51.56 |
| $R_{meas}$ [%] | 43.5 (263.9) |
| $CC_{1/2}$ | 99.0 (65.2) |
| **Data statistics for anisotropy-corrected data** | |
| Resolution range [Å] | 69.18–2.80 (2.90–2.80) |
| Unique reflections | 15 541 (626) |
| Completeness [%] | 63.60 (26.31) |
| Wilson B-factor | 34.48 |
| **Refinement statistics for anisotropy-corrected data** | |
| Reflections used in refinement | 15 541 (626) |
| Reflections used for R-free | 735 (42) |
| $R_{work}$ [%] | 22.54 (30.42) |
| $R_{free}$ [%] | 28.24 (36.73) |
| *Number of non-hydrogen atoms* | |
| Total | 5 639 |
| Macromolecules | 5 472 |
| Ligands | 136 |
| Solvent | 31 |
| Protein residues | 682 |
| RMS (bonds) [Å] | 0.001 |
| RMS (angles) [°] | 0.40 |
| Ramachandran favored [%] | 95.40 |
| Ramachandran allowed [%] | 4.60 |
| Ramachandran outliers [%] | 0.00 |
| Rotamer outliers [%] | 0.16 |
| Clashscore | 4.12 |

| | |
|---|---|
| *B-factors [Å$^2$]* | |
| Average | 48.76 |
| Macromolecules | 48.92 |
| Ligands | 47.47 |
| Solvent | 27.28 |
| Number of TLS groups | 11 |

**Extended data table 1.** Crystallographic data collection and refinement statistics for the determined structure of the KITbp–KIT123 complex (PDB ID: 9H71). Statistics for the highest-resolution shell are shown in parentheses.
